# Robust-tedana: An automated denoising pipeline for multi-echo fMRI data

**DOI:** 10.1101/2025.06.16.660050

**Authors:** Bahman Tahayori, Robert E. Smith, David N. Vaughan, Chris Tailby, Daniel A. Handwerker, Eric Y. Pierre, Graeme D. Jackson, David F. Abbott, the Australian Epilepsy Project Investigators

## Abstract

Multi-echo functional Magnetic Resonance Imaging (fMRI) data are acquired by recording image volumes at multiple echo times and can be used to improve the separation of neural activity from noise. TE-Dependent ANAlysis (tedana) is an open-source software tailored to denoising of multi-echo fMRI data. The efficacy of denoising can however be inconsistent, often necessitating manual inspection that precludes its application in large-scale studies where processing is ideally fully automated. Here, we introduce Robust-tedana, an optimised denoising pipeline that achieves adequate results at both single-subject and group level. Robust-tedana incorporates Marchenko-Pastur Principal Component Analysis (MPPCA) for effective thermal noise reduction, robust independent component analysis for stabilised signal decomposition, and a modified component classification process. We evaluated its performance on Multi-Band Multi-Echo (MBME) language-task fMRI data from the Australian Epilepsy Project (AEP) using objective measures, comparing to conventional fMRI analysis with and without multi-echo-based denoising. Experts’ manual evaluation was undertaken on a subset of these data to validate the objective measures. The proposed pipeline both mitigates the prevalence of erroneous attenuation of genuine task activation due to instability of single-subject analysis, and increases the magnitude of group-wise effects. Robust-tedana therefore facilitates advanced analysis of MBME fMRI data in an automated pipeline, including for clinical research assessment of individuals.

## 1 Introduction

Functional Magnetic Resonance Imaging (fMRI) can non-invasively map human brain function by detecting Blood Oxygenation-Level Dependent (BOLD) signal (Poldrack et al., 2011). However, fMRI data typically suffer from low Signal to Noise Ratio (SNR), where the magnitude of the BOLD signal can be difficult to distinguish from other physiological and machine effects. Multi-Echo (ME) data acquisition, where multiple volumes are recorded at different echo times (TEs) for each repetition time (TR), is a well-established method to enhance the quality of fMRI using denoising methods (Cohen, Chang, & Wang, 2021; Cohen et al., 2018; Kundu et al., 2017; Moser et al., 2025; Poser et al., 2006; Reddy et al., 2024; Steel et al., 2022; Zhao et al., 2024). These methods take advantage of the expectation that the BOLD signal manifests as modulation of the transverse magnetisation decay time constant *T* ^∗^, while nuisance physiological effects manifest differently, and the statistical features of thermal noise remain constant. To denoise the fMRI data, an image time series is decomposed into spatially independent components using Independent Component Analysis (ICA), which are then classified as either BOLD or non-BOLD depending on the echo-time dependency of their time series (Kundu et al., 2012, 2017). A drawback of multi-echo acquisition is a longer TR compared to Single Echo (SE) acquisition; Simultaneous Multi Slice (SMS) imaging techniques, also known as Multi-Band (MB) acquisitions, are often used to achieve an acceptable TR despite the presence of multiple k-space readouts per excitation (Barth et al., 2016; Xu et al., 2013).

TE Dependent ANAlysis (tedana) is an open-source tool for denoising ME and MBME fMRI data (DuPre et al., 2021), built upon the original ME-ICA framework (Kundu et al., 2012). An fMRI analysis pipeline that includes tedana typically involves multiple other software tools both upstream (eg. image realignment to correct for head motion) and downstream (eg. statistical analysis to derive a spatial map of task-related activation) of the multi-echo denoising step. The ME-ICA pipeline has been shown to outperform non-denoising pipelines in terms of activation volume and magnitude of t-statistics for both block design and event-related tasks (Gonzalez-Castillo et al., 2016). ME-ICA has further been shown effective for Multi-Band Multi-Echo (MBME) resting-state fMRI data, where the identification of more BOLD components compared to a multi-echo single-band acquisition was attributed to greater temporal resolution, facilitating identification of non-BOLD signals of higher temporal frequency (Olafsson et al., 2015).

### 1.1 Motivation

The Australian Epilepsy Project (AEP) is a national-scale project aiming to collect multi-modal imaging, cognitive and genetic data to enhance diagnosis and clinical outcome prediction for people with epilepsy. Both task-based and resting-state fMRI data are collected using a MBME sequence (Feinberg et al., 2010; Xu et al., 2013), with the intention of integrating multi-echo-based denoising into the automated image processing pipeline.

To evaluate the performance of tedana and quantify how it improves activation detection, we applied the existing implementation of tedana (v23.0.2 with default options) to the AEP language-task data (this pipeline is referred to as “Standard-tedana” pipeline throughout). In our preliminary application of Standard-tedana to these data, we noticed that at the single-subject level there were cases where Standard-tedana did not retain certain signal components that were clearly of neuronal origin. This issue has been previously reported in other studies (Cohen et al., 2017; Gonzalez-Castillo et al., 2016; Rolinski et al., 2020) and has also been acknowledged by the tedana development team. A suggested solution is to *manually* reclassify the misidentified components; eg. in (Reddy et al., 2024), the authors manually classified all components for every subject in a motor-task experiment. Reclassifying the components requires visual inspection of the tedana output by an expert for each individual subject. This humanresource-intensive task can be costly and impractical in many situations, especially for studies with a large number of subjects. Furthermore, manual classification is more difficult for resting-state fMRI data. A robust fully automatic approach is therefore highly desirable.

### 1.2 Aim

Our aim in this study was to modify the Standard-tedana denoising pipeline so that its performance would be robust at a single-subject level, suitable for integration into a fully-automated analysis pipeline.

To achieve the goal, we identified three sources of misclassifications and modified the denoising framework accordingly. 1) Estimating the number of principal components that exceed the magnitude of thermal noise is challenging; particularly given that the data input to tedana has already undergone preprocessing steps that involve interpolation of signals across voxels, which modifies the statistical properties of the data. To address this, we applied Marchenko-Pastur Principal Component Analysis (MPPCA), a method that provides an objective threshold to distinguish the signal from thermal noise, to unprocessed fMRI data (Ades-Aron et al., 2021). 2) The stochastic nature of ICA decomposition methods means a single evaluation with a particular starting seed value may result in a non-robust outcome. We therefore used a robust version of ICA (*robustica*), where many ICA iterations with different seeds are clustered together to yield a robust result (Anglada-Girotto et al., 2022). 3) Some signal independent components were inappropriately rejected as noise in the “decision tree” used by tedana to classify components. To address this issue, we identified and removed criteria that were subjectively assessed to be causing such misclassifications in data incorporating the aforementioned modifications.

The efficacy of denoising procedures within fMRI processing pipelines was evaluated based on language fMRI, where there is a strong *a priori* expectation of the relevant functional areas of the brain based on extensive prior study. Although it is desirable for these denoising methods to be applicable to resting-state fMRI data, it is difficult to objectively assess their efficacy based on their effects upon such data. We, therefore, focused here on evaluation of denoising methods using task-based fMRI data, on the premise that methods proven to have beneficial attributes under such evaluation will exhibit similar attributes when applied to resting-state data.

Activation maps derived through the three pipelines under evaluation (Robust-tedana, Standard-tedana and a pipeline that omitted any form of explicit denoising) were evaluated using two methods. We utilised the volume of supra-threshold activation within a Region of Interest (ROI) encoding the expected language activation areas in template space as an objective metric of pipeline performance (noting that all denoising methods are agnostic to the nature and timing of the task performed). We additionally validated this metric in a subset of the evaluated data by comparing its outcomes to expert evaluation of the thresholded activation maps (individuals were blinded to the denoising method used to produce each activation map).

## 2 Methods

### 2.1 Data collection

We used the Center for Magnetic Resonance Research (CMRR) Echo-Planar Imaging (EPI) sequence to collect MBME fMRI images from *n*=250 AEP participants in a 3T Siemens PrismaFit MRI scanner, with the following parameters: three echoes at TE=[15 33.25 51.5]ms, TR=0.9s, multi-band factor=4, GRAPPA factor=2, Field of View (FOV)=216 × 216 × 132mm^3^, voxel size=3 × 3 × 3mm^3^, Flip Angle (FA)=30^◦^, bandwidth=2670Hz/px, 202 volumes, anterior-posterior phase encoding direction. Acquisition included paired “blip-up/blip-down” spin-echo images to use for susceptibility distortion correction. Due to a change in project acquisition protocol, 117 subjects were acquired with a 32-channel head coil, with the other 133 subjects acquired with a 64-channel head coil. T_1_-weighted MPRAGE images with TE=2.19ms, TI=1.0s, TR=1.9s, FA=8^◦^ as well as T_2_-weighted FLAIR images with TE=392ms, TI=1.8s, TR=5s, FA=120^◦^ were acquired. For both structural images, the voxel size, FOV and GRAPPA factor were 3 × 3 × 3mm^3^, 230 × 230 × 173mm^3^ and 2, respectively. This study was approved by Austin Health human research ethics committee (HREC/60011/Austin-2019 and HREC/68372/Austin-2022).

#### • **2.1.1** Task description

All participants completed a “pseudoword rhyming” block design language fMRI task as part of the Australian Epilepsy Project (Tailby et al., 2017). During the active phase, participants determine whether visually presented pairs of pseudowords rhyme or not (e.g., “fline” vs “plyne” or “clat” vs “torb”). This task was performed silently and required grapheme-to-phoneme conversion. In the baseline phase, participants decide whether pairs of forward and backward slashes are identical (e.g., /\\\ vs \\\/), a visuospatial task. Each pair was displayed to the participant for 4.5 seconds, one above the other at the centre of the screen. The task experiment commenced with two pattern matching pairs, followed by alternating blocks of four rhyming pairs and four pattern matching pairs. The task ended with two pairs of baseline patterns, i,e. another half block to capture the downswing of the hemodynamic response function (HRF).

### 2.2 Initial assessment of Standard-tedana

As a preliminary investigation into the efficacy of the existing tedana software, we analysed the data from the first 20 subjects, applying tedana (v23.0.2 with ‘kundu’ tree) to echoes that had been preprocessed by fMRIPrep (Esteban et al., 2019) (more details are provided in Section 2.4). We observed that for some subjects, Standard-tedana erroneously removed neural activity associated with the task that manifested when the multi-echo data were simply combined naively without advanced denoising. Upon investigating this effect, we found that the detected activation depended on the arbitrary initial seed value used in the ICA step, as shown in Fig. 3. These observations motivated the pursuit of methodological solutions such that the application of multi-echo fMRI denoising would not result in erroneous inference of individualised language localisation.

### 2.3 Proposed modifications

Addressing the described issues necessitated several inter-related changes to the pipeline. These are presented as a singular cohesive set of changes, referred to as “Robust-tedana” throughout.

#### 2.3.1 Thermal noise suppression

The MPPCA method was applied to suppress thermal noise from the fMRI time series prior to preprocessing using fMRIPrep. The improvements achieved using MPPCA for fMRI data are described in (Ades-Aron et al., 2021; Henriques et al., 2023). Within tedana, thermal noise suppression via PCA truncation was then explicitly disabled, with all components proceeding to ICA.

#### 2.3.2 Robust ICA

ICA is sensitive to its initial seed value, which can lead to inconsistent results (see Fig. 3). ICA can be stabilised by performing multiple runs with different seeds, clustering the resulting components, and selecting those most consistent (Himberg & Hyvarinen, 2003). We modified the tedana code to make use of the robustica package (Anglada-Girotto et al., 2022) for this purpose.

#### 2.3.3 Modifications to selection criteria

During our initial assessment, we observed that some criteria in the default tedana classification tree removed ICA components that appeared to correspond to the task paradigm in both time and space. Therefore, we bypassed the criteria for 1) rejecting borderline components in ‘kundu’ tree(Kundu et al., 2012); 2) rejecting components with more significant voxels in magnitude than the *T* ^∗^ decay model; 3) rejecting higher dice coefficient for magnitude compared to the *T* ^∗^ decay model.

### 2.4 Processing pipelines

Data were pre-processed using fMRIPrep (Esteban et al., 2019) (ver 23.1.4) and the resulting echoes were further processed using tedana (DuPre et al., 2021) (ver 23.0.2); see Fig. 1(a). For Standard-tedana, we used the Akaike’s Information Criterion (AIC) for the Principal Component Analysis (PCA) thresholding, which is the least aggressive criterion and the default method in tedana.

**Figure 1:**
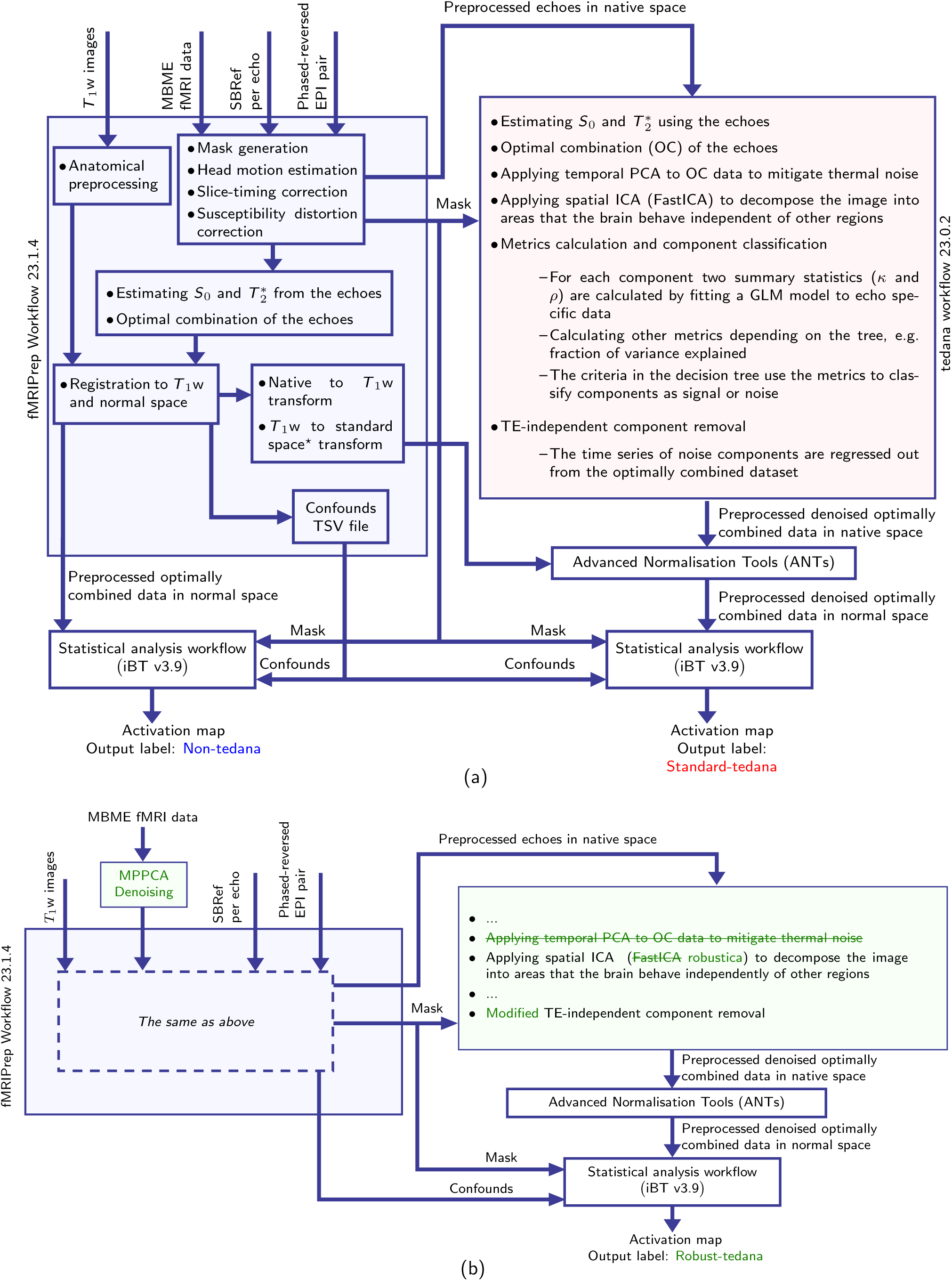
Block diagrams of the three pipelines. (a) Non-tedana and Standard-tedana pipelines to estimate the activation map for the language task; (b) Robust-tedana pipeline that contains MPPCA denoising of fMRI language data prior to any pre-processing as well as modifications to the tedana library. The same parameters are used in the statistical analysis workflow. ^*^Montreal Neurological Institute (MNI) space was used.

Our modified workflow incorporating changes to denoising is presented in Fig. 1(b). In this omnibus workflow, we first applied MPPCA denoising with a window size of 7×7×7 to mitigate thermal noise, using the dwidenoise command with double precision option from MRtrix software package (Tournier et al., 2019). As this method should usurp the thermal denoising performed in Standard-tedana based on PCA thresholding, here the PCA within tedana was instructed to preserve all variance (as its outputs were nevertheless still requisite for subsequent operations). The ‘FastICA’ algorithm from sklearn library was replaced with ‘robustica’, with the number of independent ICA runs set to 50. Furthermore, we updated the decision tree to reflect modifications described in Section 2.3.3. The output of this pipeline was labelled as “Robust-tedana”.

The preprocessed time series resulting from fMRIPrep and tedana were analysed with the Integrated Brain Analysis Toolbox for SPM, iBT version 3.9 (Abbott et al., 2024) with SPM12 (Friston et al., 2007) revision 7771. For the first level analysis, we used a General Linear Model (GLM) with identical parameters (p-value was 0.001 and False Discovery Rate (FDR) corrected p-value set to 0.05) for all three pipelines and calculated the t-score map at a single-subject level. Subsequently, the z-score map was calculated from the t-score map using spm t2z command with a number of Degrees of Freedom (DOF) determined separately for each workflow:

- For the Non-tedana pipeline the DOF was set to number of volumes included in the analysis.
- For the Standard-tedana pipeline the DOF was set to the number of components selected by the PCA step for that subject.
- For the Robust-tedana pipeline the DOF was set to the minimum of the rank determined by the MPPCA step and the number of components given by ‘robustica’.

### 2.5 Language mask creation

Evaluation of the performance of the three methods in an automated and objective manner necessitated definition of a region of interest (ROI) in which activity was expected for the task performed. A language mask was generated from a subset of 60 subjects, whom were excluded from all subsequent evaluations. For analyses utilising this mask to be unbiased with respect to all methods under evaluation, we split this cohort into 3 sets of 20, with each set contributing data pre-processed using one of the three methods. From these data, a language mask was computed using a Random-Effects (RFX) analysis in SPM; this is shown in red in Fig. 2.

**Figure 2:**
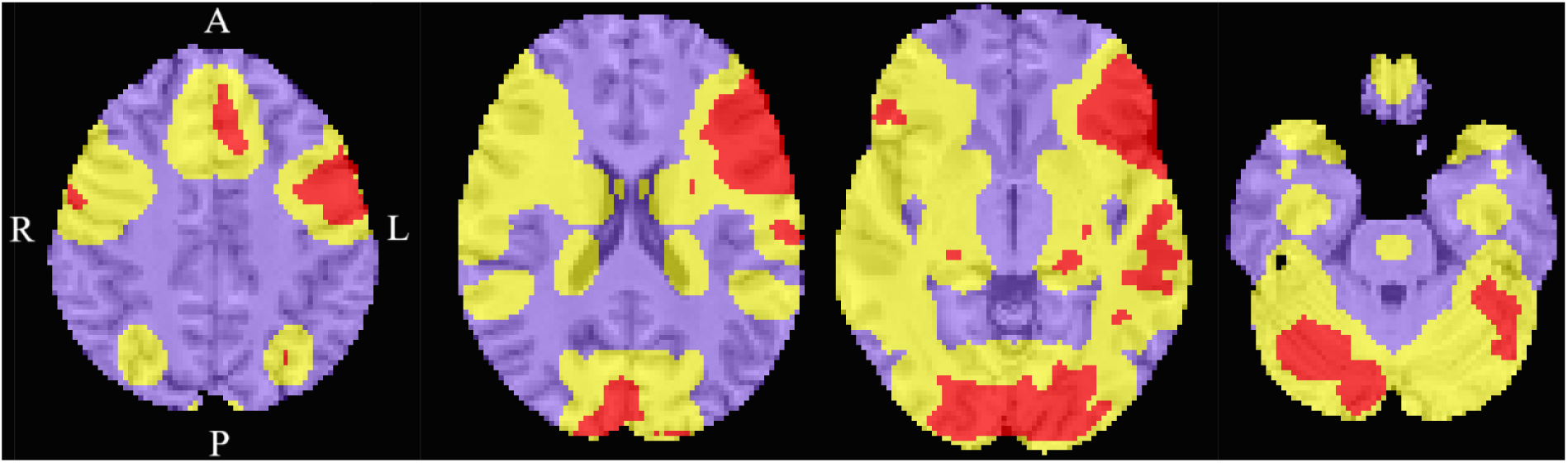
The derived template space region of interest corresponding to areas of expected language activation. The mask derived from random effects analysis is shown in red; ‘ignored’ area shown in yellow corresponds to a dilated version of this mask for exclusion from subsequent ROC analysis. The purple section is the ‘non-active’ ROI.

While this mask was sufficient for the calculation of activation volumes within expected language regions, further manipulation was necessary to facilitate classification of supra-threshold voxels as “true positive” vs. “false positive”. In a quantitative comparison of preprocessing methods, neither a supra-threshold voxel immediately adjacent to this mask, nor a supra-threshold voxel appearing in a homologous region in the opposing hemisphere (perhaps due to non-standard language lateralisation), could be confidently interpreted as erroneous detection of activation. We therefore generated an explicit ‘ignored’ area using the following series of steps: dilated the language mask using an 18mm spherical kernel (notably larger than the 8 mm smoothing performed in the statistical analysis step); computed the union between this mask and its reflection through the mid-sagittal plane; taking the complement of this mask. The result of this computation is shown in yellow in Fig. 2. The rest of the brain was considered ‘non-active’ and characterised by purple in Fig. 2.

### 2.6 Criteria for comparing pipelines

To systematically compare the performance of the three pipelines, we used four metrics that are computed independently per subject and denoising method.

1. Activation volume: This is the number of supra-threshold voxels (*p* = 0.001) within the language ROI (red in Fig. 2).
2. Mean z-score: The mean value of the z-statistic map within the language ROI.
3. The Area Under the Curve (AUC): We generated Receiver Operating Characteristic (ROC) curves using the z-statistic values within the language (red in Fig. 2) and non-active area (purple in Fig. 2) regions of the template brain. We used the perfcurve command in MATLAB.
4. Temporal Signal-to-Noise Ratio (tSNR): We calculated voxelwise tSNR across the brain producing a histogram and mean tSNR per subject and denoising method. This measure is commonly used to assess denoising methods (Beckers et al., 2023; Cohen, Jagra, et al., 2021; Gilmore et al., 2022; Vizioli et al., 2021).

### 2.7 Experts’ evaluation

We engaged in manual evaluation of task activation maps for two reasons: to ascertain whether the differences in outcomes of fMRI processing caused by changes to denoising procedures were consequential for the current primary use case of such data, and to validate the suitability of our data-driven metrics of language activation map quality when applied in an automated fashion to the larger cohort. The evaluation was performed by two clinicians from our centre with expertise in language network identification.

#### 2.7.1 Cohort selection

30 subjects were selected from the cohort for expert assessment, with all 30 subjects assessed by both experts to facilitate investigation of inter-rater variance. To ensure that this cohort would include the full breadth of discriminatory behaviour between denoising methods, we applied a convex hull algorithm to activation volume data and included within this subset those subjects on this hull (Fig.4(a)-(f)). We manually included additional subjects where at least one method yielded a very small activation volume, and finally filled the cohort with a random selection of additional subjects.

**Figure 3:**
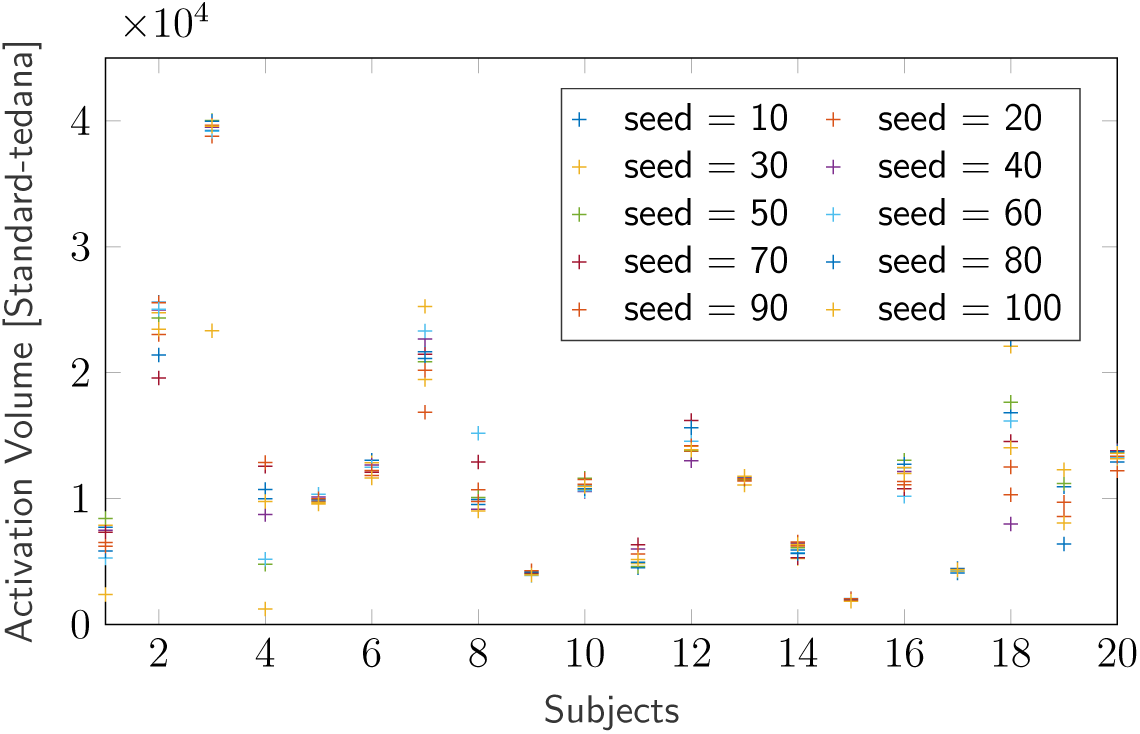
The seed dependency of the Standard-tedana pipeline. Each mark shows the total volume of supra-threshold voxels throughout the brain when executing the Standard-tedana pipeline with the same input data, varying only the ICA seed value.

**Figure 4:**
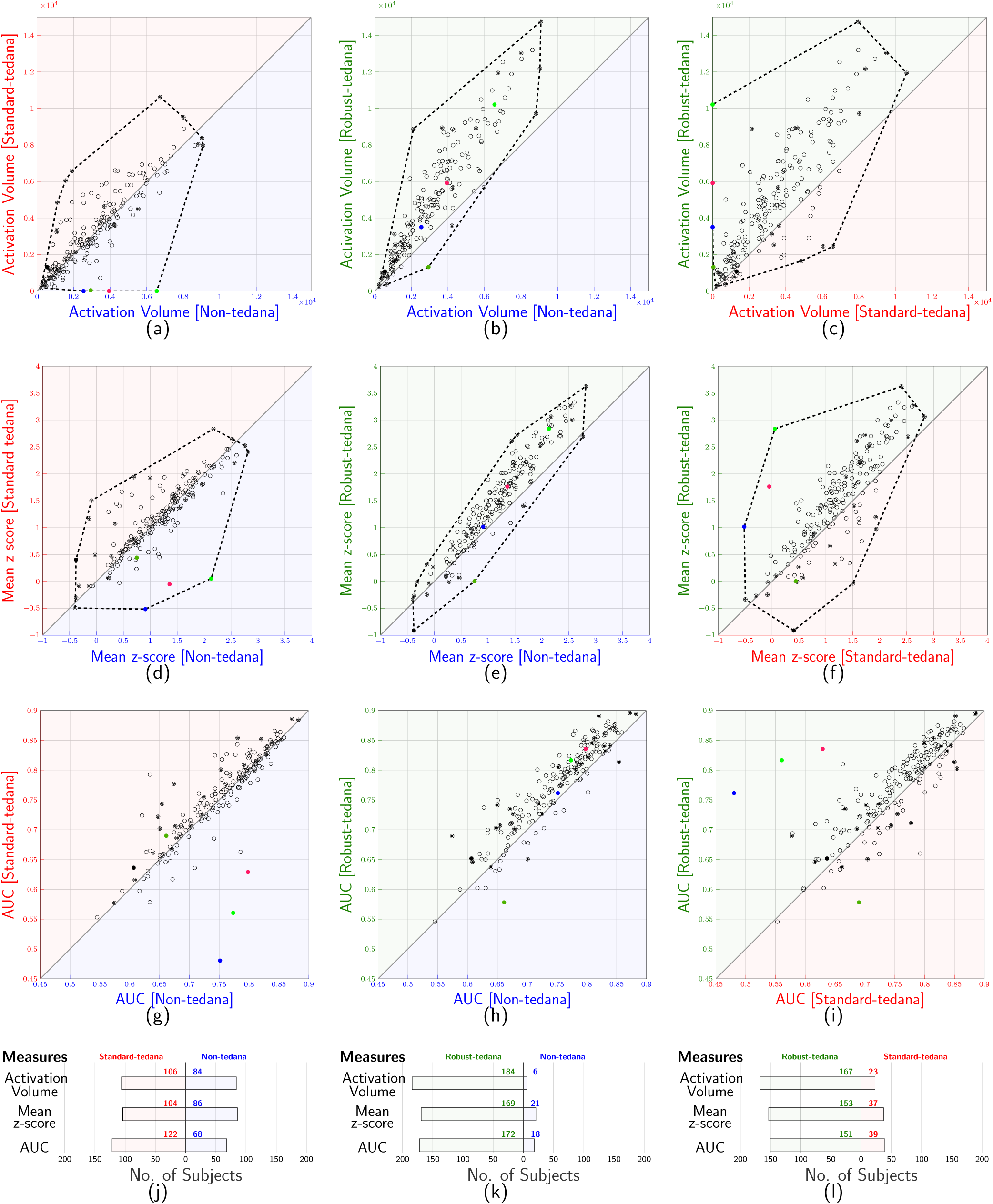
Pairwise comparison of the three pipelines in terms of: (a)-(c) activation volume within the task ROI; (d)-(f) mean z-score within the task ROI; and (g)-(i) the Area Under the Curve (AUC) within the brain for each subject. Each axis corresponds to the results derived from one specific pipeline. Each circle represents data from one subject. The 30 subjects selected for expert evaluation are distinguished by filled circles. Dashed lines show the convex hulls that were used to identify extreme cases for inclusion in the expert evaluation cohort. In (j)-(l), a summary of the result is presented. The coloured points show how outliers moved when Robust-tedana pipeline was used.

#### 2.7.2 Details of manual assessment

We designed a three-way comparison by setting up three separate two-way comparisons; 1) Non-tedana vs. Standard-tedana 2) Non-tedana vs. Robust-tedana and 3) Standard-tedana vs. Robust-tedana. Activation maps were superimposed on the subject’s brain in light-box format. For each pair-wise comparison, activation maps from two pipelines were presented side-by-side on one page, with randomisation of order of the two methods. Clinicians were blind to the method that created each activation map.

In each pairwise comparison, we asked the experts to answer the following questions:

1. For each activation map individually, is the map *of adequate quality to be interpretable*? For this question a simple binary response was requested for each of the two maps.
2. Of the two language activation maps presented on the page, *which is physiologically more plausible*? Clinicians responded on a 5 point scale positioned horizontally underneath the side by side activation maps, and ran 2, 1, 0, 1, 2. A response of 2 or 1 on the left indicated a preference for the left image, whereas a response of 1 or 2 on the right indicated a preference for the right image. The middle option (0) denoted that both methods produced equally plausible (or non-plausible) results, and the remaining responses (1 or 2) indicated that one method was “superior” or “much superior”. For clarity, an example page of the document is shown in the supplementary material (Fig. S1).

#### 2.7.3 Validation of activation volume measure

Pairwise comparison of the activation volume measure with experts’ evaluation was performed using the following metric:

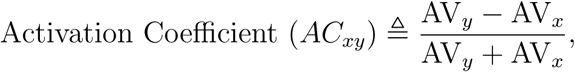

where *x* and *y* are identifiers of specific denoising pipelines, and AV_i_ represents the activation volume detected by pipeline *i*. This measure is zero-centred, such that *AC*_*xy*_ = 0.0 corresponds to equivalent activation volume between pipelines, with values greater/smaller than 0.0 indicating greater activation volume from pipeline *y*/*x*, respectively. Validation of this measure was performed by correlating it against the ordinal 5-point-scale scores registered by the expert clinicians.

## 3 Results

### 3.1 Seed dependency of Standard-tedana

In the initial assessment of Standard-tedana pipeline using 20 subjects, we quantified the activation volume across the whole brain for 10 different tedana ICA initial seed values using the seed option in tedana; see Fig 3. The detected activation volume for several subjects altered substantially with the seed value, with extreme cases exhibiting changes of over an order of magnitude for different tedana seeds.

### 3.2 Comparison of pipelines

The pairwise results of the three pipelines in terms of activation volume and mean z-score within the task ROI are presented in Figs. 4(a)-(f). The AUCs are shown in Figs. 4(g)-(i).

In Figs. 4(a)-(i), each circle represents one subject from the cohort. Convex hull analysis (dashed lines in Figs. 4(a)-(f)) identified 16 of the extreme cases for inclusion in the cohort for expert assessment, all of which are shown as filled circles.

The summary in Fig. 4(j) suggests that the Standard-tedana pipeline has a small tendency to yield more extensive language activation than the Non-tedana pipeline. There were however subjects that showed zero or negligible detected activity when analysed with the Standard-tedana pipeline, contrary to the evidence of substantial activation with the simpler Non-tedana pipeline; four of these subjects are given unique colour indicators in Fig. 4(a), with these subject identifiers propagated to other sub-figures of Fig. 4.

Fig. 4(k) shows that the Robust-tedana pipeline exhibits a more substantial improvement to activation detection over the Non-tedana pipeline than does the Standard-tedana pipeline (Fig. 4(j)) in terms of all measures. More importantly, in contrast to the Standard-tedana pipeline, there are no subjects with evidence of task-related activity via the activation volume measure when analysed with the Non-tedana pipeline that is entirely attenuated through application of the Robust-tedana pipeline, Fig. 4(j)–4(k). Comparison of the Standard-tedana and Robust-tedana pipelines is summarised in Fig. 4(l), which indicates that for 80% of the subjects Robust-tedana has a higher performance compared to the Standard-tedana pipeline.

The mean Receiver Operating Characteristic (ROC) curves for all three pipelines are presented in Fig. 5. The mean AUC of the Robust-tedana pipeline is significantly higher than Non-tedana and Standard-tedana pipelines with *p* ≪ 0.001, with no significant difference between the mean AUC of Non-tedana and Standard-tedana pipelines (*p* = .12).

**Figure 5:**
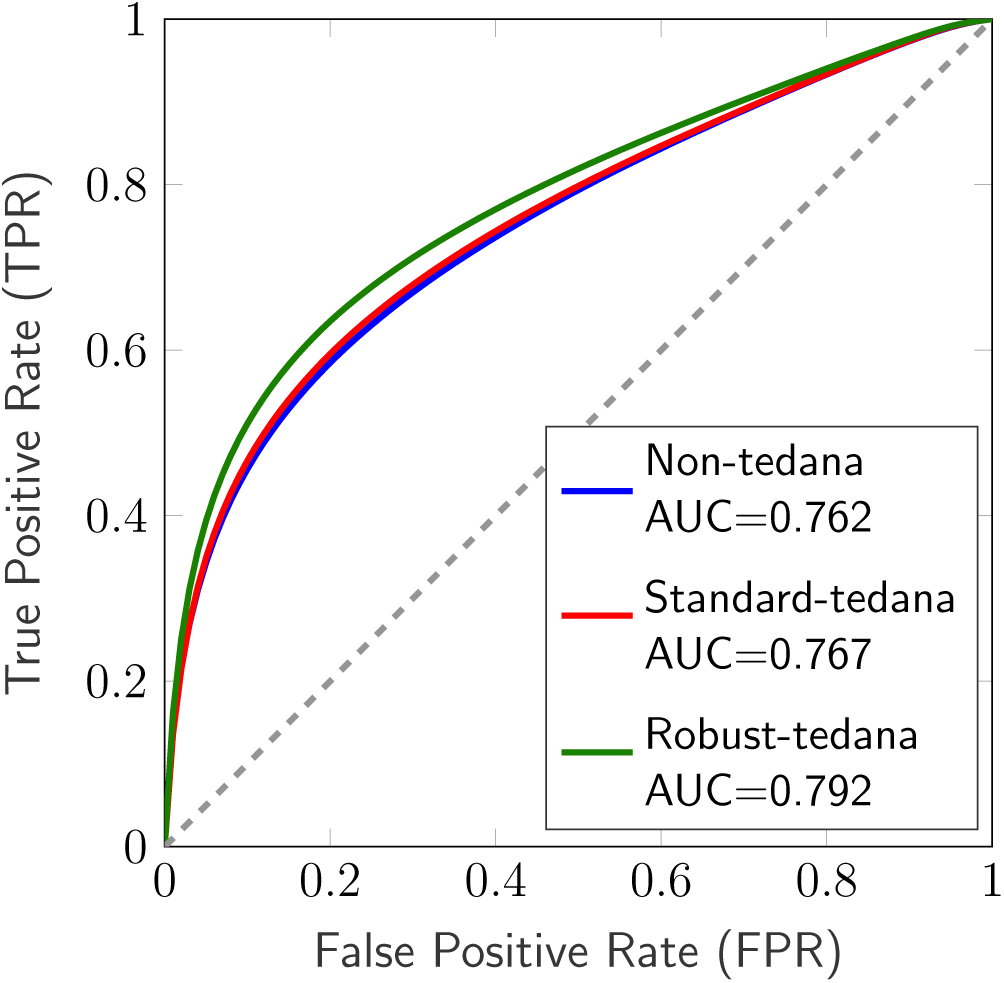
The mean ROC for all three methods. Robust-tedana has a significantly higher mean AUC than Non-tedana and Standard-tedana with *p* ≪ 0.001. There was no significant difference between the mean AUC of Non-tedana and Standard-tedana pipelines.

Fig. 6 shows the detected activation for three examples selected from cases where Standard-tedana performed relatively poorly (but not as poorly as the complete failures coloured in Fig. 4). This figure illustrates how Robust-tedana improved the activation detection for these subjects.

**Figure 6:**
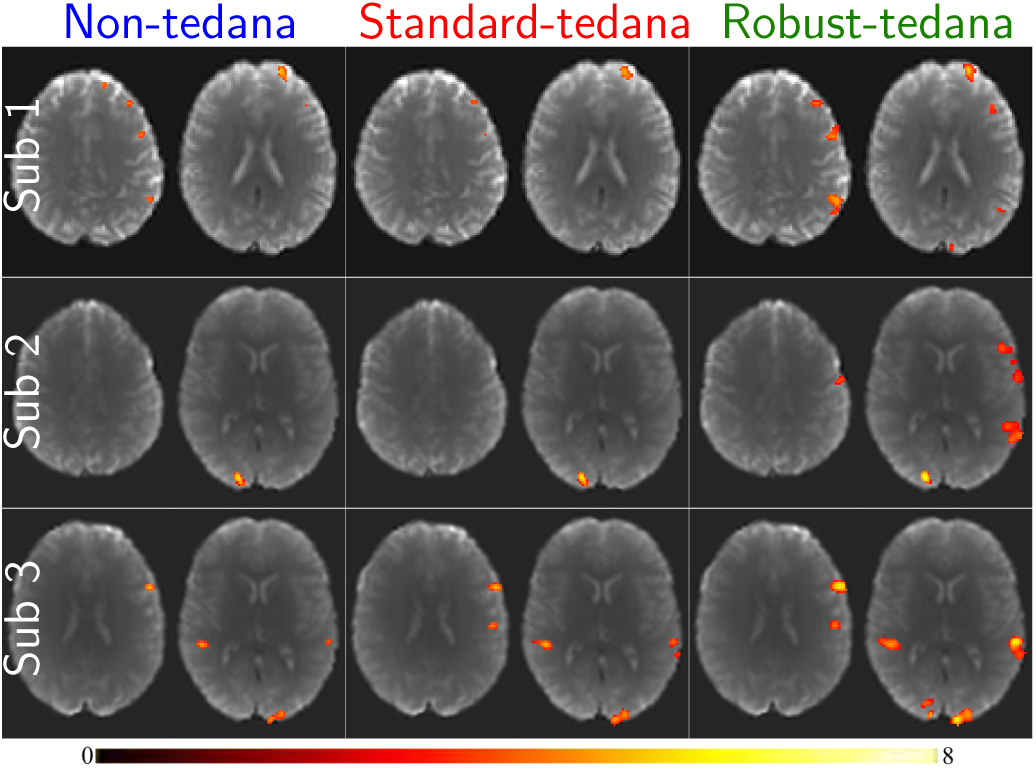
Detected activation in a language task for three subjects using Non-tedana, Standardtedana and Robust-tedana pipelines. Each column represents a pipeline, while each row corresponds to a subject. Two axial slices are shown for each subject. The detected activation volumes for Subject 1 are (AV-Non-tedana, AV-Standard-tedana, AV-Robust-tedana)=(504, 277, 1026). For Subjects 2 and 3 the activation volumes by each method are (400, 346, 989) and (908, 1133, 2160), respectively.

### 3.3 Experts’ evaluation result

Fig. 7(a)-(c) shows scatter plots of activation volumes in a manner akin to Fig. 4, for only the subset of 30 subjects that underwent manual evaluation, but additionally with the results of those manual evaluations overlaid. Each square represents a subject, colour coded evaluation according to the 5-scale option explained in Section 2.7, with the upper half of the square representing Clinician 1 and the lower half representing Clinician 2. The horizontal and vertical half-lines indicate clinician responses to the question of clinical interpretability of the activation maps by the methods labelled on the x-axis and y-axis, respectively. Unlike Fig. 4, here logarithmic scales are employed to better distinguish all individual subjects across the full spectra of activation volumes.

**Figure 7:**
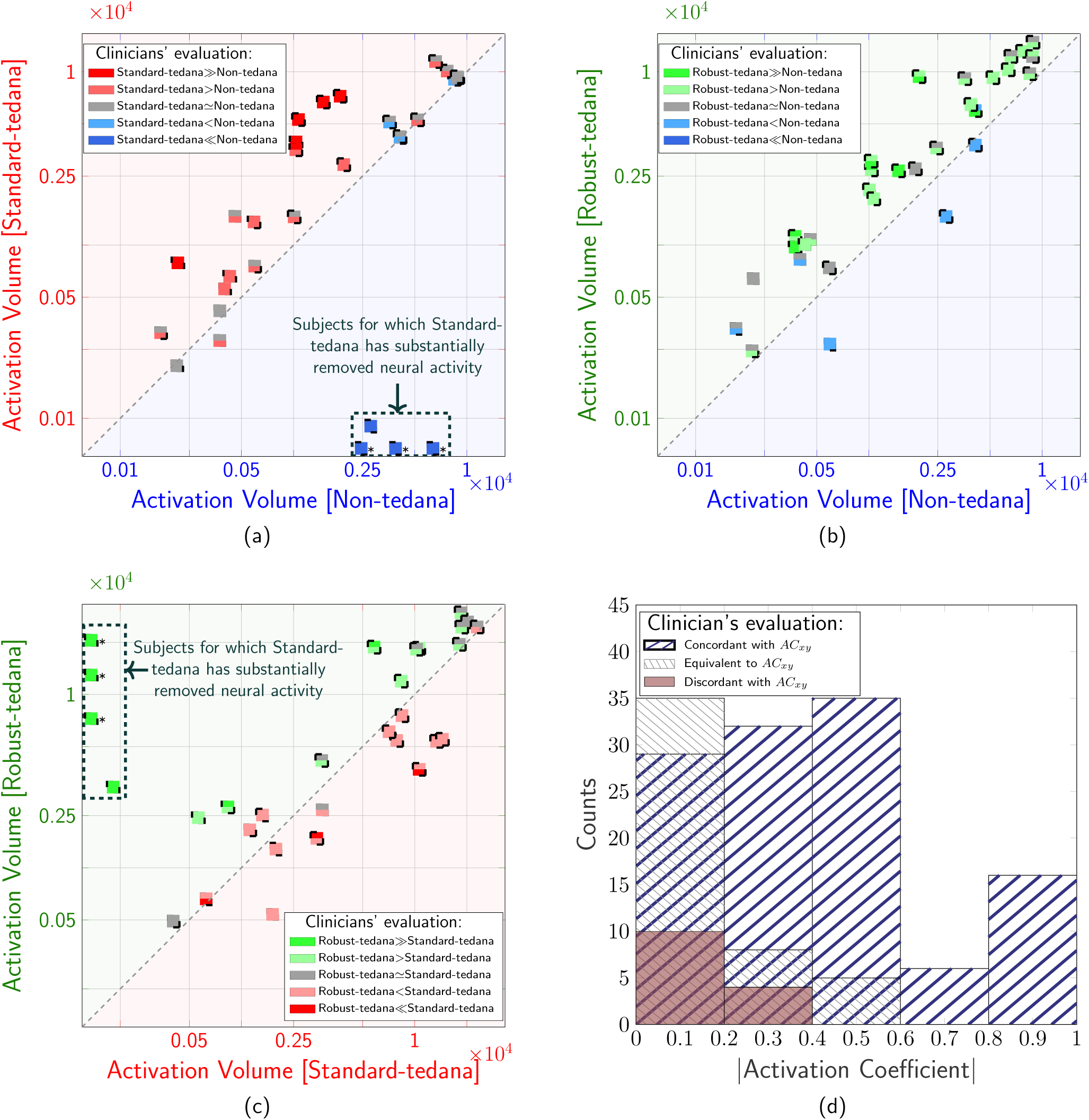
Experts’ evaluation for 30 subjects in a pairwise comparison superimposed on activation volume plots. Pipelines under comparison are: (a) Standard-tedana versus Non-tedana; (b) Robust-tedana versus Non-tedana; (c) Robust-tedana versus Standard-tedana. Each square represents one of the subjects distinguished by a filled circle in Fig. 4(a)-(c). For each square, the upper half represents Clinician 1 and the lower half represents Clinician 2. The horizontal and vertical half-lines indicate each clinician response to the question of clinical interpretability of the presented activation map by the method labelled on the x-axis and y-axis respectively. The three subjects marked with asterisks showed no activity when the Standard-tedana pipeline was used. To include these subjects in the logarithmic plots, their data was added manually. (d) shows a histogram to summarise all of the concordance ratings in (a)-(c) versus the absolute value of Activation Coefficient. “Concordant” or “Discordant” are defined as when the equality line and an expert comparison are in agreement or disagreement, respectively; “Equivalent” represents comparisons scored equal by the clinicians. Interpretation: (a) demonstrates that standard tedana is most often superior compared to analyses without tedana (most points are above the equality line), however, standard tedana is not robust as there are subjects with little to no activity only when tedana is applied (these subjects are made clear by the red dashed rectangle); (b) demonstrates that Robust-tedana is most often superior to Non-tedana; (c) illustrates that Robust-tedana and Standard-tedana perform similarly in most cases (largely even distribution of subjects around the line of equivalence), Standard-tedana sometime fails (subjects in the red rectangle), whereas Robust-tedana always succeeded; (d) illustrates good concordance between expert evaluation and objective activation volume measurement. This is also evident in (a)-(c) as most points have the same colour as the background colour of the half-planes separated by the equality line. The only points where there is some discordance occurred close to equivalence on the activation volume measure (in the vicinity of *AC*_*xy*_ = 0).

Clinician assessments in Figs. 7(a)-(c) largely align with the equality lines, supporting activation volume as a reasonable quantitative marker of pipeline performance. Fig. 7(d) shows the concordance between the equality line classifier and clinicians as a function of *AC*_*xy*_. Here, “Concordant” or “Discordant” indicate agreement or disagreement between the equality line and expert ratings, respectively. “Equivalent” represents comparisons scored equal by clinicians. This histogram shows that the discordance is more likely in the vicinity of *AC*_*xy*_ = 0, while performance improves as *AC*_*xy*_ deviates from this value. There was discordance in only 14 comparisons (8%) whereas 118 comparisons (66%) were concordant.

### 3.4 Temporal Signal-to-Noise Ratio (tSNR)

The tSNR for the denoising pipelines is shown in Fig. 8. Two additional pipelines are shown over and above the three pipelines described elsewhere in the article. The first involved extracting the Non-tedana pipeline only data from the second echo, prior to fitting the GLM, thus reflecting the loss of specificity from no longer performing optimal combination of data across multiple echoes, compared to the Non-tedana pipeline. The second involved applying MPPCA denoising prior to the Non-tedana pipeline, but not utilising either version of tedana, therefore reflecting the magnitude of influence of specifically the MPPCA method relative to the Non-tedana pipeline. Fig. 8(a) shows the histogram of tSNR values across the brain volumes of all subjects. Fig. 8(b) presents box plots encoding, for each pipeline, the distribution of the mean tSNR computed for each subject. It quantifies the incremental enhancement of the tSNR at each step. The maximum improvement is achieved with the Robust-tedana pipeline when all modifications proposed in Section 2 are in place.

**Figure 8:**
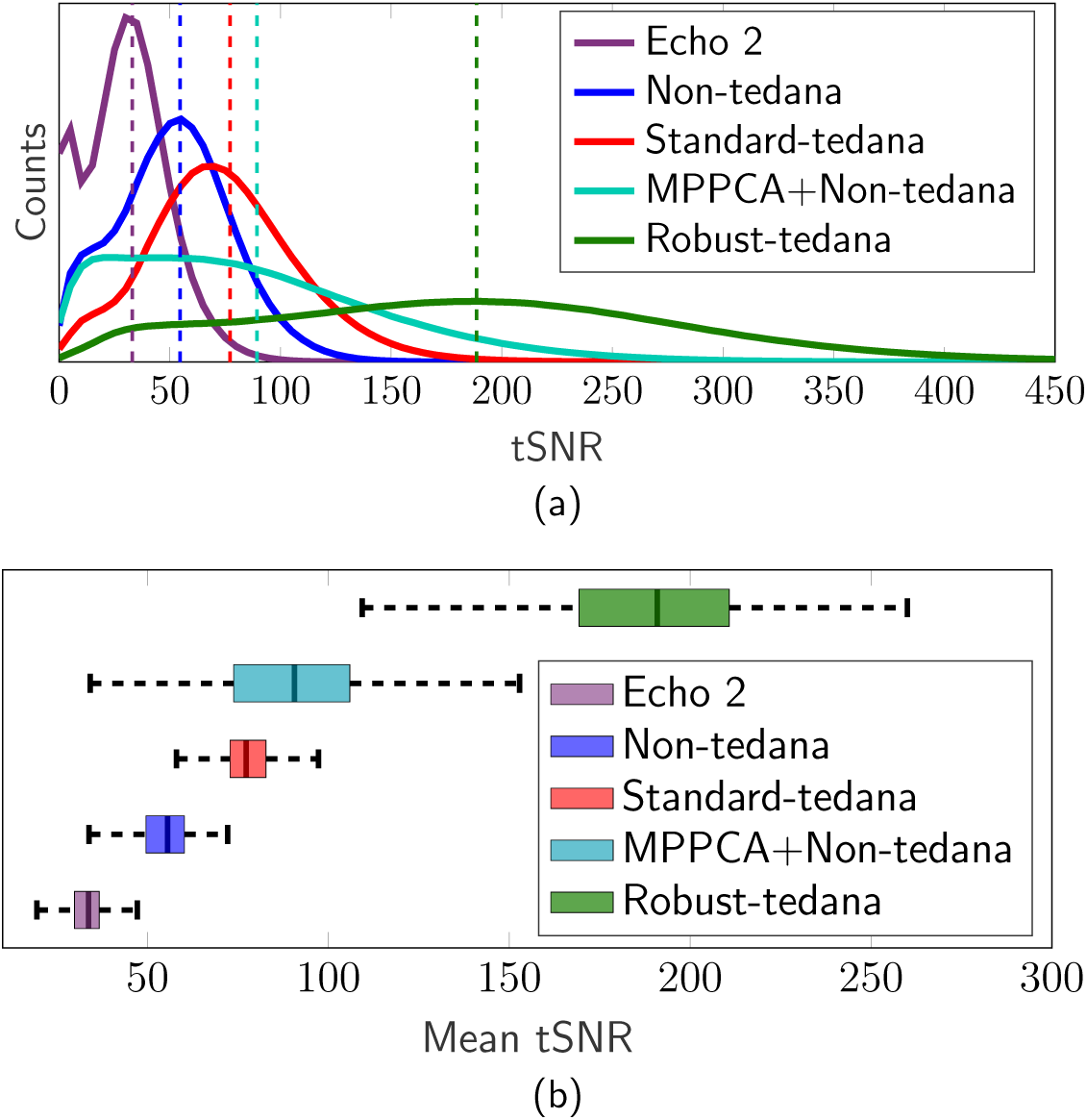
Whole-brain statistics of temporal-SNR (tSNR) for different pipelines. “Echo 2”: Modification of Non-tedana pipeline where only data from the second echo was provided as input to the GLM. “MPPCA+Non-tedana”: Modification of Non-tedana pipeline where MPPCA was applied prior to fMRIPrep, but tedana was not utilised. (a) Histograms of tSNR through the brain volume aggregated over all subjects for each pipeline; vertical dashed lines show the mean tSNR and (b) Distribution of mean tSNR across subjects for each pipeline.

## 4 Discussion

We proposed a denoising pipeline for multi-echo fMRI data that uses Marchenko-Pastur PCA denoising, fMRIPrep, and a modified version of the tedana software; see Fig. 1(b). We analysed a large cohort of task-based MBME fMRI data from the Australian Epilepsy Project (AEP) with three different pipelines (Fig. 1). We showed that while the inclusion of the tedana software into the analysis pipeline yielded greater sensitivity to group effects, in several subjects it attenuated signals clearly originating from neuronal activity. This could be highly deleterious to clinical interpretation if integrated into a large-scale fully automated pipeline. We presented an alternative processing pipeline, incorporating multiple modifications that target the root cause of this instability at the single subject level. This substantially enhances the viability of multi-echo fMRI acquisitions across a broad range of use cases, including individualised clinical research assessments.

### 4.1 Validity of comparative measure

For this work, we required a data-driven quantification of the efficacy of the pipelines under consideration, which could be performed across a large cohort, as we needed to identify individual cases where a pipeline might yield deleterious results. We therefore sought to validate an automated and objective metric that would avoid the extensive manual labour of human experts. We showed, in a subgroup of the cohort, there was strong concordance between expert assessment of language activation maps with a simple measure of activation volume within a language ROI. This was observed in a held-out subset of the cohort using the same task paradigm and acquisition protocol. These findings justify the use of this measure throughout our work as a summary metric of the sensitivity of different preprocessing pipelines to *a priori* expected task-related activity.

### 4.2 On thermal denoising

A key modification of the pipeline that we propose here is not only the mechanism by which thermal noise mitigation is performed, but the stage within the pipeline at which that denoising is performed. The tedana software includes a PCA step, where one of multiple heuristic thresholding techniques may be employed to withhold noisy components from the subsequent ICA, where classification of BOLD-like vs. non-BOLD-like components occurs. However, the fact that this step is performed subsequent to preprocessing using a tool such as fMRIPrep, which invariably results in interpolation of input image intensities to produce the preprocessed data, makes objective separation of thermal noise from genuine signal extremely difficult. Use of the Moving Average PCA (MAPCA) approach here can mitigate this effect, by reducing the extent to which a single input image intensity may modify multiple preprocessed image intensities, thereby breaking expectations of independence of observations. However, determination of an appropriate number of components to retain is problematic. Indeed, the tedana software provides multiple PCA thresholding criteria, due to the ambiguity of this step.

PCA-based thermal denoising has arguably a stronger history in the domain of diffusion MRI data, where the image contrast of interest is generated through substantial signal attenuation, often resulting in image data very close to the thermal noise floor (Veraart et al., 2016). There are two key aspects of state-of-the-art developments in this domain that are directly applicable to the construction of a robust multi-echo fMRI pipeline:

1. It is well-established that these methods Dowdle et al., 2023; Moeller et al., 2021; Olesen et al., 2023; Vizioli et al., 2021; Zhu et al., 2022 are ineffective for thermal denoising if they are applied after other image processing steps that involve interpolation of data– e.g. gradient non-linearity distortion correction, *k*-space zero-padding, slice timing correction, motion correction, resulting in non-independence of intensity samples (Ades-Aron et al., 2021). Given that some of these manipulations may be applied by the scanner itself, researchers interested in replicating this pipeline are encouraged to ensure that the data they regularly obtain from their own scanner hardware are not compromised.
2. The Marchenko-Pastur law provides mathematical rigour to the expected manifestation of random noise in the PCA eigenspectrum, and therefore an objective and unambiguous mechanism by which to optimally suppress the presence of thermal noise through filtering of that eigenspectrum.

### 4.3 Robust ICA

The ‘robustica’ Python library stabilises ICA by running ‘FastICA’ multiple times and deriving a consensus decomposition, (Anglada-Girotto et al., 2022). The computational burden is therefore increased compared to running just a single ICA within the Standard-tedana pipeline. In our analysis, tedana in the Standard-tedana pipeline (1 ICA run) took about 4 minutes, whereas the Robust-tedana pipeline with 50 ICA runs took roughly 27 minutes on the same system (Intel Xeon Gold 6448H) using single-core single-thread with 16 GB of memory.

A convergence analysis of robust ICA clustering based on the number of runs is presented in the supplementary material to justify the selection of 50 runs for all analyses, see Fig. S2. Furthermore, we created a 2-d projection map of the components using T-distributed Stochastic Neighbour Embedding (TSNE) so that the user can visualise the output of the clustering process. This graph is added to the tedana html report. An example of a projection graph is shown in supplementary material, see Fig. S3.

### 4.4 Limitations

Given that MPPCA is a sliding window approach, the signal rank, and therefore effective DOF, is nonstationary. Ideally, this should be accounted for in any fMRI analysis pipelines utilising any denoising approach based on a sliding window. Failure to do so will result in erroneously inflated test statistics and therefore a false positive rate greater than that intended. Given that the suite of software tools in use here are not yet adequately equipped to deal with this confound, we performed a conservative post-hoc correction for the Robust-tedana pipeline, wherein the DOF to transform from *t*-values to *z* -scores was set to the minimum of the component count yielded by robustica and the signal rank estimated by MPPCA for the patch centred at each image voxel.

### 4.5 Applicability to resting-state fMRI

A major prospective application of multi-echo fMRI denoising is in the study of brain networks at rest (Cohen et al., 2017; Lynch et al., 2020; Pilmeyer et al., 2023). It has been shown that the use of MBME data with ICA denoising can increase functional connectivity (Cohen, Jagra, et al., 2021). Our work here focused exclusively on the use of task-based fMRI specifically because validation of the proposed pipeline necessitated data with a strong *a priori* expectation of where activation should and should not occur. It is however expected that fMRI paradigms without block task designs will nevertheless be a major application for our modified Robust-tedana pipeline.

## 5 Conclusion

We proposed Robust-tedana, a denoising pipeline for multi-echo fMRI datasets that is based on MPPCA denoising and modification of the tedana open-source library. We used task fMRI data from the Australian Epilepsy Project (AEP) to demonstrate how the pipeline mitigates erroneous suppression of task-related activation in individuals, critical for clinical research assessment. Clinicians’ input was used to validate that ROI activation volume is a suitable objective measure of the performance of pipelines. Robust-tedana showed robust performance at the individual subject level. The pipeline can be used for denoising large cohorts where automated pipelines are desirable. We chose to validate the algorithm’s performance on task fMRI data because it is more straightforward to assess expected activation patterns. The pipeline is fully automatic, and is applicable regardless of the presence or absence of a block design fMRI task. We expect the greatest utility of the denoising approach will be in application to multi-band multi-echo resting-state fMRI data, where noise-removal is an even more critical element of the analysis.

## Data and code availability

Any requests for access to the data used in this project should be directed to the Australian Epilepsy Project (a formal data access request can be lodged at https://www.epilepsyproject.org.au/research/ access-to-aep-data). All components of the Robust-tedana pipeline are available in open-source software packages. The modified decision tree file (robust tedana 2025.json) and its associated flowchart can be found at https://figshare.com/articles/code/component selection decision trees in tedana/25251433/ 3.

## Author contributions

B.T: Theoretical development, framework conceptualisation, methodology, coding and running simulations, data analysis and manuscript drafting. R.E.S.: Theoretical development, framework conceptualisation, critical insights and discussions to refine the model and manuscript, coding, manuscript drafting and manuscript review. D.N.V.: Conceptualisation, methodology, investigation, resources, writing - review & editing, project administration, funding acquisition and clinical evaluation of the pipeline. C.T.: Conceptualisation, methodology, investigation, resources, data curation, writing - original draft, writing - review & editing, Visualisation, Supervision, project administration, funding acquisition and clinical evaluation of the pipeline. D.A.H.: Model development, critical insights and discussions for refining the model (particularly related to the tedana package), interpretation of the results and manuscript review. E.Y.P.: Experimental design, technical support and manuscript review. G.D.J.: Conceptualisation, methodology, investigation, resources, writing - review & editing, project administration and funding acquisition. D.F.A.: Conceptualisation, methodology, software, investigation, resources, data curation, writing - original draft, writing - review & editing, visualisation, supervision, project administration, funding acquisition, theoretical development, critical insights and discussions to refine the model and manuscript and revising the manuscript for intellectual content.

## Declaration of competing interest

The authors declare no competing interests.

## Acknowledgements

We acknowledge the facilities and scientific and technical assistance of the National Imaging Facility, a National Collaborative Research Infrastructure Strategy (NCRIS) capability, at The Florey Institute of Neuroscience and Mental Health. We also acknowledge the strong support from the Victorian Government and in particular the funding from the Operational Infrastructure Support Grant, and support from the Victorian Biomedical Imaging Capability (VBIC). The Australian Epilepsy Project received funding from the Australian Government under the Medical Research Future Fund (Frontier Health and Medical Research Program - Grant Numbers MRFF75908 and RFRHPSI000008) and the Victoria State Government (Victorian-led Frontier Health and Medical Research Program). We acknowledge receipt of software for the multi-echo multiband fMRI sequence from the University of Minnesota Center for Magnetic Resonance Research. This research was supported by The University of Melbourne’s Research Computing Services and the Petascale Campus Initiative.

The authors gratefully acknowledge Dr. Eneko Uruñuela of the University of Calgary for his contribution in integrating the TSNE plot into the auto-generated HTML report in tedana v25.0.0.

## Notes

### Competing Interest Statement

The authors have declared no competing interest.

## References

Abbott, D., Capon, A., Tailby, C., Tahayori, B., Bryant, M., Vaughan, D., Jackson, G., & for the Australian Epilepsy Project Investigators. (2024). The Integrated Brain Analysis Toolbox for SPM (iBT). Proceedings of the Organization for Human Brain Mapping Annual Meeting (OHBM). https://archive.aievolution.com/2024/hbm2401/Abstracts/viewAbs?abs=2320

Ades-Aron, B., Lemberskiy, G., Veraart, J., Golfinos, J., Fieremans, E., Novikov, D. S., & Shepherd, T. (2021). Improved task-based functional MRI language mapping in patients with brain tumors through Marchenko-Pastur principal component analysis denoising. Radiology, 298 (2), 365–373. 10.1148/radiol.2020200822

Anglada-Girotto, M., Miravet-Verde, S., Serrano, L., & Head, S. A. (2022). robustica: customizable robust independent component analysis. BMC Bioinformatics, 23 (1), 519. 10.1186/s12859-022-05043-9

Barth, M., Breuer, F., Koopmans, P. J., Norris, D. G., & Poser, B. A. (2016). Simultaneous multislice (SMS) imaging techniques. Magnetic Resonance in Medicine, 75 (1), 63–81. 10.1002/mrm.25897

Beckers, A. B., Drenthen, G. S., Jansen, J. F., Backes, W. H., Poser, B. A., & Keszthelyi, D. (2023). Comparing the efficacy of data-driven denoising methods for a multi-echo fMRI acquisition at 7T. Neuroimage, 280, 120361. 10.1016/j.neuroimage.2023.120361

Cohen, A. D., Chang, C., & Wang, Y. (2021). Using multiband multi-echo imaging to improve the robustness and repeatability of co-activation pattern analysis for dynamic functional connectivity. NeuroImage, 243, 118555. 10.1016/j.neuroimage.2021.118555

Cohen, A. D., Jagra, A. S., Yang, B., Fernandez, B., Banerjee, S., & Wang, Y. (2021). Detecting task functional MRI activation using the multiband multiecho (MBME) echo-planar imaging (EPI) sequence. Journal of Magnetic Resonance Imaging, 53 (5), 1366–1374. 10.1002/ jmri.27448

Cohen, A. D., Nencka, A. S., Lebel, R. M., & Wang, Y. (2017). Multiband multi-echo imaging of simultaneous oxygenation and flow timeseries for resting state connectivity. PloS ONE, 12 (3), e0169253. 10.1371/journal.pone.0169253

Cohen, A. D., Nencka, A. S., & Wang, Y. (2018). Multiband multi-echo simultaneous ASL/BOLD for task-induced functional MRI. PLoS ONE, 13 (e019042). 10.1371/journal.pone.0190427

Dowdle, L. T., Vizioli, L., Moeller, S., Aķcakaya, M., Olman, C., Ghose, G., Yacoub, E., & Uğurbil, K. (2023). Evaluating increases in sensitivity from NORDIC for diverse fMRI acquisition strategies. NeuroImage, 270, 119949. 10.1016/j.neuroimage.2023.119949

DuPre, E., Salo, T., Ahmed, Z., Bandettini, P. A., Bottenhorn, K. L., Caballero-Gaudes, C., Dowdle, L. T., Gonzalez-Castillo, J., Heunis, S., Kundu, P., et al. (2021). TE-dependent analysis of multiecho fMRI with tedana. Journal of Open Source Software, 6 (66), 3669. 10.21105/joss.03669

Esteban, O., Markiewicz, C. J., Blair, R. W., Moodie, C. A., Isik, A. I., Erramuzpe, A., Kent, J. D., Goncalves, M., DuPre, E., Snyder, M., et al. (2019). fMRIPrep: a robust preprocessing pipeline for functional MRI. Nature Methods, 16 (1), 111–116. 10.1038/s41592-018-0235-4

Feinberg, D. A., Moeller, S., Smith, S. M., Auerbach, E., Ramanna, S., Glasser, M. F., Miller, K. L., Ugurbil, K., & Yacoub, E. (2010). Multiplexed echo planar imaging for sub-second whole brain fMRI and fast diffusion imaging. PloS ONE, 5 (12), e15710. 10.1371/journal.pone.0015710

Friston, K. J., Ashburner, J., Kiebel, S. J., Nichols, T., & Penny, W. (2007). Statistical parametric mapping. 10.1016/B978-0-12-372560-8.X5000-1

Gilmore, A. W., Agron, A. M., Gonźalez-Araya, E. I., Gotts, S. J., & Martin, A. (2022). A comparison of single-and multi-echo processing of functional MRI data during overt autobiographical recall. Frontiers in Neuroscience, 16, 854387. 10.3389/fnins.2022.854387

Gonzalez-Castillo, J., Panwar, P., Buchanan, L. C., Caballero-Gaudes, C., Handwerker, D. A., Jangraw, D. C., Zachariou, V., Inati, S., Roopchansingh, V., Derbyshire, J. A., & Bandettini, P. A. (2016). Evaluation of multi-echo ICA denoising for task based fMRI studies: Block designs, rapid eventrelated designs, and cardiac-gated fMRI. NeuroImage, 141, 452–468. 10.1016/j.neuroimage.2016.07.049

Henriques, R. N., Ianuş, A., Novello, L., Jovicich, J., Jespersen, S. N., & Shemesh, N. (2023). Efficient PCA denoising of spatially correlated redundant MRI data. Imaging Neuroscience, 1, 1–26. 10.1162/imag_a_00049

Himberg, J., & Hyvarinen, A. (2003). Icasso: software for investigating the reliability of ICA estimates by clustering and visualization. 2003 *IEEE XIII Workshop on Neural Networks for Signal Processing (IEEE Cat. No. 03TH8718)*, 259–268. 10.1109/NNSP.2003.1318025

Kundu, P., Inati, S. J., Evans, J. W., Luh, W.-M., & Bandettini, P. A. (2012). Differentiating BOLD and non-BOLD signals in fMRI time series using multi-echo EPI. NeuroImage, 60 (3), 1759–1770. 10.1016/j.neuroimage.2011.12.028

Kundu, P., Voon, V., Balchandani, P., Lombardo, M. V., Poser, B. A., & Bandettini, P. A. (2017). Multiecho fMRI: A review of applications in fMRI denoising and analysis of BOLD signals. NeuroImage, 154, 59–80. 10.1016/j.neuroimage.2017.03.033

Lynch, C. J., Power, J. D., Scult, M. A., Dubin, M., Gunning, F. M., & Liston, C. (2020). Rapid precision functional mapping of individuals using multi-echo fMRI. Cell Reports, 33 (12). 10.1016/j.celrep.2020.108540

Moeller, S., Pisharady, P. K., Ramanna, S., Lenglet, C., Wu, X., Dowdle, L., Yacoub, E., Uğurbil, K., & Aķcakaya, M. (2021). NOise reduction with DIstribution Corrected (NORDIC) PCA in dMRI with complex-valued parameter-free locally low-rank processing. Neuroimage, 226, 117539. 10.1016/j.neuroimage.2020.117539

Moser, J., Nelson, S. M., Koirala, S., Madison, T. J., Labonte, A. K., Carrasco, C. M., Feczko, E., Moore, L. A., Lundquist, J. T., Weldon, K. B., et al. (2025). Multi-echo acquisition and thermal denoising advances precision functional imaging. Imaging Neuroscience, 3, imag_a_00426. 10.1162/imag_a_00426

Olafsson, V., Kundu, P., Wong, E. C., Bandettini, P. A., & Liu, T. T. (2015). Enhanced identification of BOLD-like components with multi-echo simultaneous multi-slice (MESMS) fMRI and multi-echo ICA. NeuroImage, 112, 43–51. 10.1016/j.neuroimage.2015.02.052

Olesen, J. L., Ianus, A., Østergaard, L., Shemesh, N., & Jespersen, S. N. (2023). Tensor denoising of multidimensional MRI data. Magnetic Resonance in Medicine, 89 (3), 1160–1172. 10.1002/mrm.29478

Pilmeyer, J., Hadjigeorgiou, G., Lamerichs, R., Breeuwer, M., Aldenkamp, A. P., & Zinger, S. (2023). Spatial and Temporal Consistency of Brain Networks for different Multi-Echo fMRI Combination Methods. IEEE Access. 10.1109/ACCESS.2023.3324183

Poldrack, R. A., Mumford, J. A., & Nichols, T. E. (2011). Handbook of functional MRI data analysis. Cambridge University Press. 10.1017/CBO9780511895029

Poser, B. A., Versluis, M. J., Hoogduin, J. M., & Norris, D. G. (2006). BOLD contrast sensitivity enhancement and artifact reduction with multiecho EPI: parallel-acquired inhomogeneity-desensitized fMRI. Magnetic Resonance in Medicine, 55 (6), 1227–1235. 10.1002/mrm.20900

Reddy, N. A., Zvolanek, K. M., Moia, S., Caballero-Gaudes, C., & Bright, M. G. (2024). Denoising task-correlated head motion from motor-task fMRI data with multi-echo ICA. Imaging Neuroscience, 2, 1–30. 10.1162/imag_a_00057

Rolinski, R., You, X., Gonzalez-Castillo, J., Norato, G., Reynolds, R. C., Inati, S. K., & Theodore, W. H. (2020). Language lateralization from task-based and resting state functional MRI in patients with epilepsy. Human Brain Mapping, 41 (11), 3133–3146. 10.1002/hbm.25003

Steel, A., Garcia, B. D., Silson, E. H., & Robertson, C. E. (2022). Evaluating the efficacy of multi-echo ICA denoising on model-based fMRI. NeuroImage, 264. 10.1016/j.neuroimage.2022.119723

Tailby, C., Abbott, D. F., & Jackson, G. D. (2017). The diminishing dominance of the dominant hemisphere: Language fMRI in focal epilepsy. NeuroImage: Clinical, 14, 141–150. 10.1016/j.nicl.2017.01.011

Tournier, J.-D., Smith, R., Raffelt, D., Tabbara, R., Dhollander, T., Pietsch, M., Christiaens, D., Jeurissen, B., Yeh, C.-H., & Connelly, A. (2019). MRtrix3: A fast, flexible and open software framework for medical image processing and visualisation. NeuroImage, 202, 116137. 10.1016/j.neuroimage.2019.116137

Veraart, J., Fieremans, E., & Novikov, D. S. (2016). Diffusion MRI noise mapping using random matrix theory. Magnetic Resonance in Medicine, 76 (5), 1582–1593. 10.1002/mrm.26059

Vizioli, L., Moeller, S., Dowdle, L., Aķcakaya, M., De Martino, F., Yacoub, E., & Uğurbil, K. (2021). Lowering the thermal noise barrier in functional brain mapping with magnetic resonance imaging. Nature Communications, 12 (1), 5181. 10.1038/s41467-021-25431-8

Xu, J., Moeller, S., Auerbach, E. J., Strupp, J., Smith, S. M., Feinberg, D. A., Yacoub, E., & Ŭgurbil, K. (2013). Evaluation of slice accelerations using multiband echo planar imaging at 3 T. NeuroImage, 83, 991–1001. 10.1016/j.neuroimage.2013.07.055

Zhao, L. S., Raithel, C. U., Tisdall, M. D., Detre, J. A., & Gottfried, J. A. (2024). Leveraging multi-echo EPI to enhance BOLD sensitivity in task-based olfactory fMRI. Imaging Neuroscience, 2, 1–15. 10.1162/imag_a_00362

Zhu, W., Ma, X., Zhu, X.-H., Ugurbil, K., Chen, W., & Wu, X. (2022). Denoise functional magnetic resonance imaging with random matrix theory based principal component analysis. IEEE Transactions on Biomedical Engineering, 69 (11), 3377–3388. 10.1109/TBME.2022.3168592

