## Supplementary material for "Robust-tedana: An automated denoising pipeline for multi-echo fMRI data"

*Correspondence: {bahman.tahayori,robert.smith,david.abbott}@florey.edu.au

### 1 Experts' evaluation

One page of the document used for methods evaluation, which was provided to the clinicians, is shown in Fig. S1.

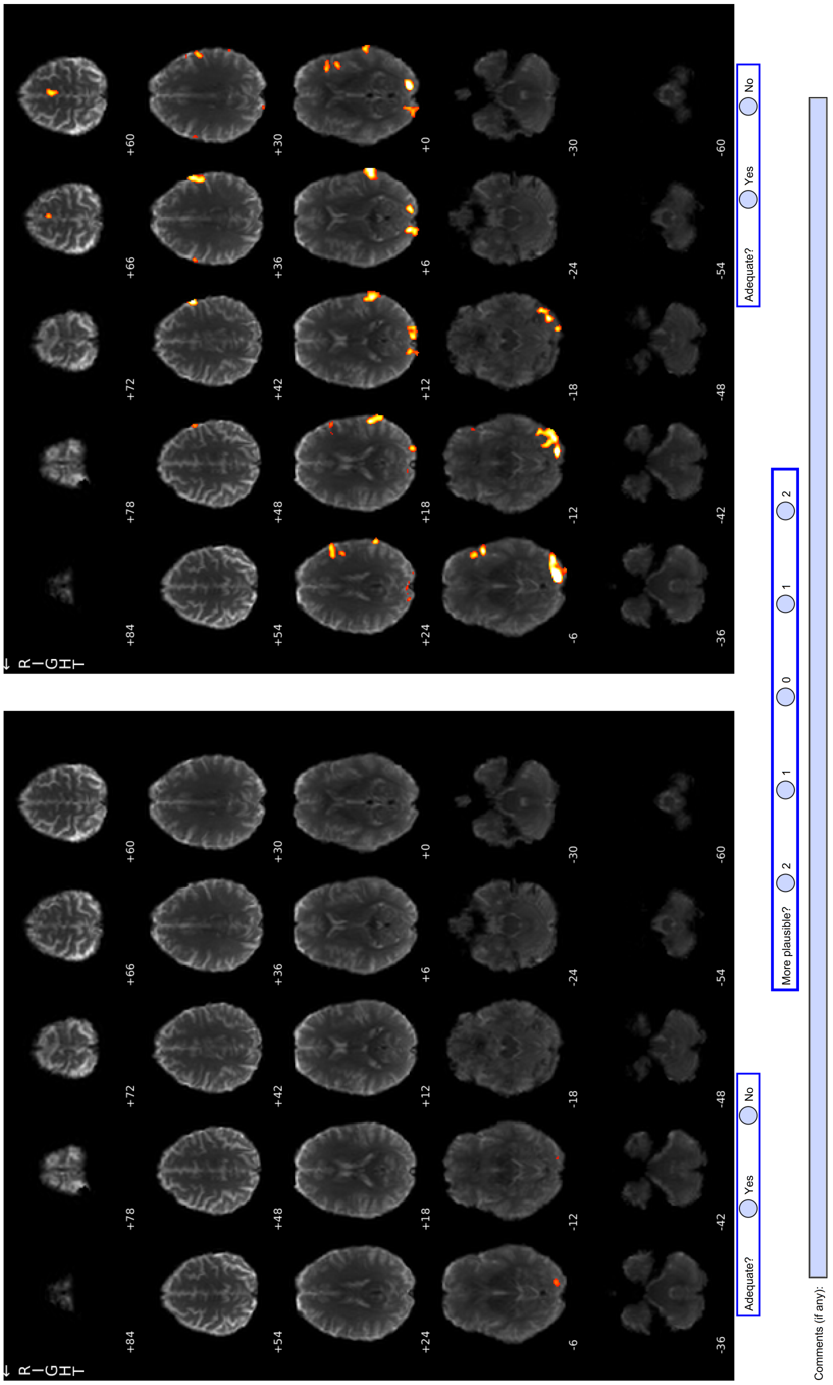


Figure S1: Example page from the document used in the methods evaluation, as presented to clinicians.

### 2 Requisite number of ICA runs for RobustICA convergence

Utilising only a very small number of ICA runs in order to reduce computational burden risks obtaining an outcome that differs substantially from what would otherwise be obtained more robustly from a much larger number of runs (a so-called "bronze standard"). We sought to choose a suitable number of runs for our analyses with some guarantee regarding the maximal deviation of the outcome from this bronze standard. Our evaluation of this effect involved a random selection of 30 subjects from the cohort. For each subject, the bronze standard outcome was generated based on execution of 'robutsica' with 200 runs. For each candidate number of ICA runs $k$, we generated 100 trials based on a random selection of $k$ ICA runs from this set of 200. For each of those trials, we applied the Density-Based Spatial Clustering of Applications with Noise (DBSCAN) clustering procedure (Ester et al., 1996), the same procedure used in 'robutsica', to the $k$ selected ICA runs, and computed the cosine similarity between that outcome and the bronze standard. We then selected the *minimum* of these cosine similarities across the 100 trials for that $k$, as this represents the worst case scenario deviation from the ideal outcome.

Figure S2 shows, for each candidate number of ICA runs $k$, the distribution across those 30 subjects of this minimum cosine similarity from the bronze standard. A non-uniform sampling of $k$ values was used to provide higher resolution at lower $k$ where convergence behaviour changes most rapidly. The results show that with as few as 24 runs, a minimum similarity above 0.80 is achievable. In this paper, we opted for a more conservative threshold, targeting a minimum similarity of 0.90; the resulting requirement of 50 runs (marked by the green dashed line in Fig. S2) was used for all analyses in this manuscript.

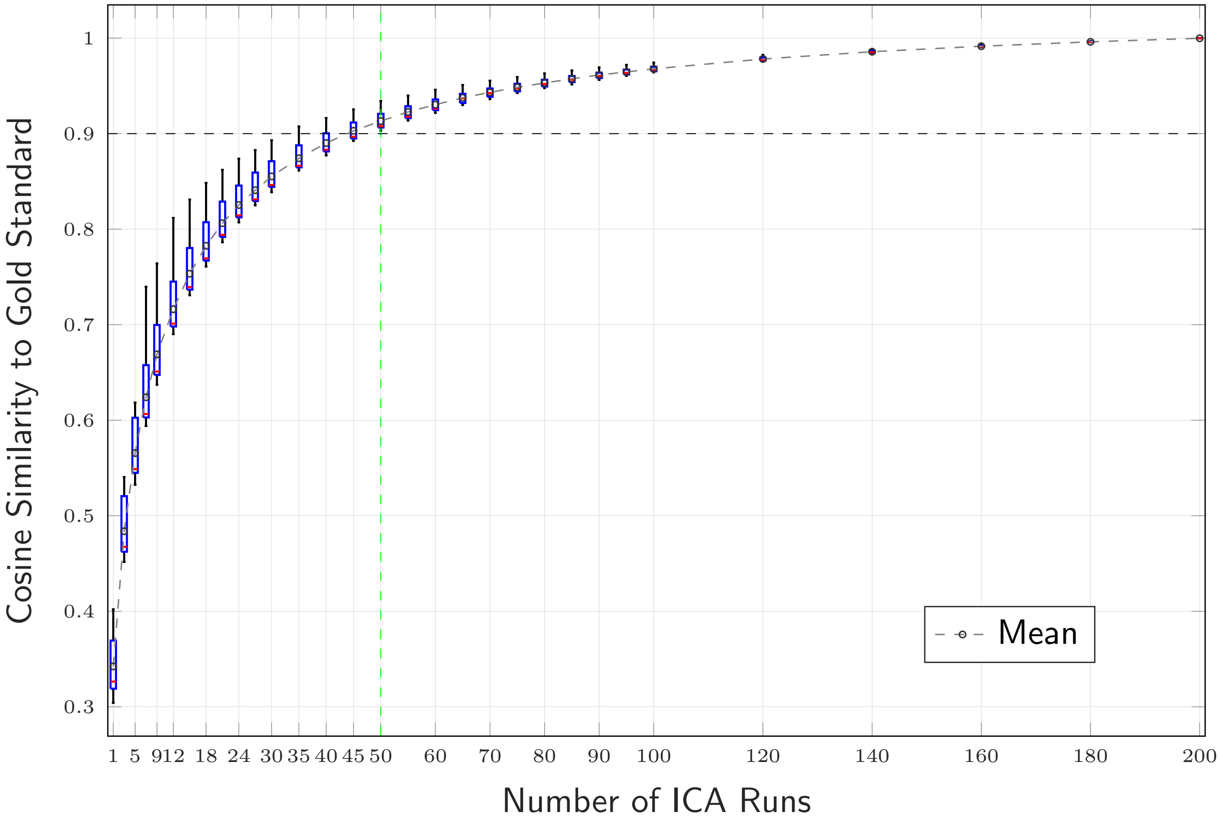


Figure S2: Convergence of robust ICA mixing matrices as a function of the number of ICA runs. For each number of runs, 100 random subsets of ICA runs were selected to compute a robust mixing matrix that is minimally similar to the bronze standard (estimated from all 200 runs).

### 3 Projection map of the clustering process

To visualise the clustering behaviour in Robust-tedana, we generated a 2D t-distributed Stochastic Neighbor Embedding (TSNE) projection (Van der Maaten & Hinton, 2008) of the component estimates across multiple ICA runs, see Fig S3. Each point in the plot represents a component from one run. Outliers that were not assigned to any cluster by the DBSCAN algorithm are shown with a red cross. The projection map illustrates the consistency of component clustering across runs.

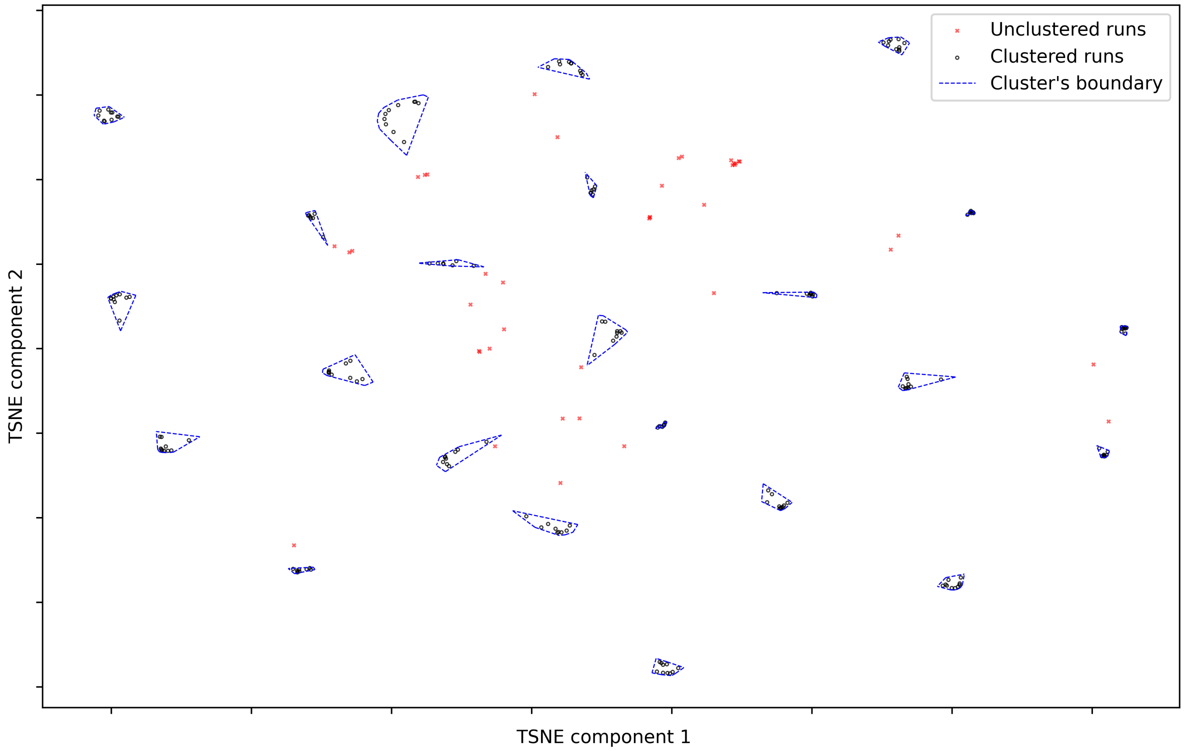


Figure S3: The TSNE projection map of the clustering process for robustica. From tedana software version 25.0.0, the same projection approach is applied to the decomposition of the input data, with the resulting figure embedded in the auto-generated HTML report.

### 4 Australian Epilepsy Project (AEP) investigator list

Australian Epilepsy Project (AEP) investigator list with Contributor Roles Taxonomy (CRediT) author statement relevant for this manuscript is shown in Table S1.

Table S1: List of AEP contributors, their affiliations, roles, and CRediT contributions.

| Name & ORCID | Primary Location | Role | CRediT |
|---|---|---|---|
| Graeme D. Jackson, MD 0000-0002-7917-5326 | The Florey Institute of Neuroscience and Mental Health | Chief Investigator | Conceptualisation, Methodology, Investigation, Resources, Writing - Review & Editing, Supervision, Project Administration, Funding Acquisition |
| David F. Abbott, PhD 0000-0002-7259-8238 | The Florey Institute of Neuroscience and Mental Health | Informatics Lead | Conceptualisation, Methodology, Software, Investigation, Resources, Data Curation, Writing - Original Draft, Writing - Review & Editing, Visualisation, Supervision, Project Administration, Funding Acquisition |
| Zanfina Ademi, PhD 0000-0002-0625-3522 | Monash University | Health Economics Lead | Conceptualisation, Funding Acquisition |
| Subhaga Amarasekara | The Florey Institute of Neuroscience and Mental Health | Product Lead | Resources, Project Administration |

(Continued on next page)

*(Continued from previous page)*

| Name & ORCID | Primary Location | Role | CRediT |
|---|---|---|---|
| Amanda Anderson | The Florey Institute of Neuroscience and Mental Health | Lived Experience Ambassador and Participant Lead | Investigation, Resources, Funding Acquisition |
| Rachel Hughes | The Florey Institute of Neuroscience and Mental Health | Clinical Research Coordinator | Investigation, Resources |
| Donna Hutchison | The Florey Institute of Neuroscience and Mental Health | Executive Lead | Project Administration |
| Patrick Kwan, MD 0000-0001-7310-276X | Monash University | Outcomes Lead | Conceptualisation, Resources, Funding Acquisition |
| Paul Lightfoot | The Florey Institute of Neuroscience and Mental Health | Operations Lead | Investigation, Project Administration |
| Saul Mullen, MD, PhD 0000-0003-1224-4101 | The University of Melbourne | Protocol Development Lead (2019–2021) | Conceptualisation, Methodology, Funding Acquisition |
| Karen L. Oliver, PhD 0000-0001-5188-6153 | The University of Melbourne | Genetics Lead | Conceptualisation, Funding Acquisition |
| Heath R. Pardoe, PhD 0000-0002-0123-2167 | The Florey Institute of Neuroscience and Mental Health | Science Operations Lead | Investigation, Resources, Writing - Review & Editing, Project Administration |

*(Continued on next page)*

*(Continued from previous page)*

| **Name & ORCID** | **Primary Location** | **Role** | **CRediT** |
|---|---|---|---|
| Mangor Pedersen, PhD 0000-0002-9199-1916 | Auckland University of Technology | Artificial Intelligence Lead | Conceptualisation, Methodology, Funding Acquisition |
| Chris Tailby, PhD 0000-0002-1320-5924 | The Florey Institute of Neuroscience and Mental Health | Neuropsychology Lead | Conceptualisation, Methodology, Investigation, Resources, Data Curation, Writing - Original Draft, Writing - Review & Editing, Visualisation, Supervision, Project Administration, Funding Acquisition |
| David N. Vaughan, MD, PhD 0000-0002-6225-7739 | The Florey Institute of Neuroscience and Mental Health | Imaging Lead | Conceptualisation, Methodology, Investigation, Resources, Writing - Review & Editing, Project Administration, Funding Acquisition |
| Anton De Weger 0009-0006-7478-361X | The Florey Institute of Neuroscience and Mental Health | Digital and Technology Lead | Software, Resources, Data Curation |

### References

Ester, M., Kriegel, H.-P., Sander, J., Xu, X., et al. (1996). A density-based algorithm for discovering clusters in large spatial databases with noise. *kdd*, *96*(34), 226–231. https://dl.acm.org/doi/10.5555/3001460.3001507

Van der Maaten, L., & Hinton, G. (2008). Visualizing data using t-SNE. *Journal of machine learning research*, *9*(11). https://www.jmlr.org/papers/volume9/vandermaaten08a/vandermaaten08a.pdf
